# Structural basis for mismatch surveillance by CRISPR/Cas9

**DOI:** 10.1101/2021.09.14.460224

**Authors:** Jack P. K. Bravo, Mu-Sen Liu, Ryan S. McCool, Kyungseok Jung, Kenneth A. Johnson, David W. Taylor

## Abstract

The widespread use of CRISPR/Cas9 as a programmable genome editing tool has been hindered by off-target DNA cleavage (Cong et al., 2013; Doudna, 2020; Fu et al., 2013; Jinek et al., 2013). While analysis of such off-target editing events have enabled the development of Cas9 variants with greater discrimination against mismatches (Chen et al., 2017; Kleinstiver et al., 2016; Slaymaker et al., 2016), the underlying molecular mechanisms by which Cas9 rejects or accepts mismatches are poorly understood (Kim et al., 2019; Liu et al., 2020; Slaymaker and Gaudelli, 2021). Here, we used kinetic analysis to guide cryo-EM structure determination of Cas9 at different stages of mismatch surveillance. We observe a distinct, previously undescribed linear conformation of the duplex formed between the guide RNA (gRNA) and DNA target strand (TS), that occurs in the presence of PAM-distal mismatches, preventing Cas9 activation. The canonical kinked gRNA:TS duplex is a prerequisite for Cas9 activation, acting as a structural scaffold to facilitate Cas9 conformational rearrangements necessary for DNA cleavage. We observe that highly tolerated PAM-distal mismatches achieve this kinked conformation through stabilization of a distorted duplex conformation via a flexible loop in the RuvC domain. Our results provide molecular insights into the underlying structural mechanisms that may facilitate off-target cleavage by Cas9 and provides a molecular blueprint for the design of next-generation high fidelity Cas9 variants that selectively reduce off-target DNA cleavage while retaining efficient cleavage of on-target DNA.

## Main

For therapeutic applications of CRISPR/Cas9, off-target DNA cleavage must be minimized (Cong et al., 2013; Fu et al., 2013; Jinek et al., 2013). While a variety of high-fidelity Cas9 variants with improved mismatch discrimination have been developed (Chen et al., 2017; Slaymaker and Gaudelli, 2021), their enhanced specificity comes at the cost of severely reduced on-target DNA cleavage rates (Kim et al., 2020; Liu et al., 2020). While mismatches induce alternative Cas9 conformations (Singh et al., 2018; Sternberg et al., 2015), the structures used to guide rational redesign of such variants were bound to on-target DNA and inactive conformations (Anders et al., 2014; Jiang et al., 2016). In order to understand the underlying molecular mechanisms underlying off-target recognition, we used kinetic analysis to guide sample preparation for cryo-electron microscopy (cryo-EM) to provide structural snapshots of Cas9 pre-cleavage activation intermediates in the presence of various DNA:RNA mismatches.

We measured the rates of target strand (TS) cleavage by Cas9 in the presence of contiguous triple nucleotide mismatches at different positions along the gRNA:TS duplex (**Extended Data Fig. 1a**). Compared to rapid on-target cleavage (1.0 s^-1^) the well-characterized PAM-distal 18-20 MM (three mismatches 18-20 bp distal from the PAM) caused a ∼40-fold reduction in rate. Other mismatches (6-8 MM, 9-11 MM and 15-17 MM) resulted in much slower rates of cleavage, plateauing at ∼20% product formed after 2 hours (**Extended Data Fig. 1b**).

Unexpectedly, the 12-14 MM allowed Cas9 activation but with rates ∼10-fold slower than the 18-20 MM. While Cas9 cleavage of 12-14 MM and 18-20 MM-containing DNA are both significantly slower than on-target, more than 80% of either substrate is cleaved within an hour of incubation with Cas9. This time-frame is highly relevant for genome editing applications, which typically occurs on the time-scale of days to weeks (Ran et al., 2013).

### Cryo-EM reveals mismatch-induced Cas9 intermediates

To understand the structural basis for Cas9 activation, we vitrified Cas9 with 12-14 MM DNA after a 5-minute of reaction, where only ∼25% of DNA has been cleaved. We determined a cryo-EM structure at a global resolution of 3.6 Å (**Fig. 1a, Extended Data Fig. 2**). The TS-cleaving endonuclease HNH was not observed due to HNH flexibility prior to activation (Dagdas et al., 2017; Zhu et al., 2019). Surprisingly, the distal end of the gRNA:TS duplex was in a linear conformation relative to the PAM-proximal TS:NTS duplex, a state that is dissimilar from previously-determined on-target DNA-bound Cas9 structures, which depict a kinked duplex (∼70°) (Jiang et al., 2016; Zhu et al., 2019), but reminiscent of early R-loop formation intermediates (Cofsky et al., 2021).

**Fig. 1.**
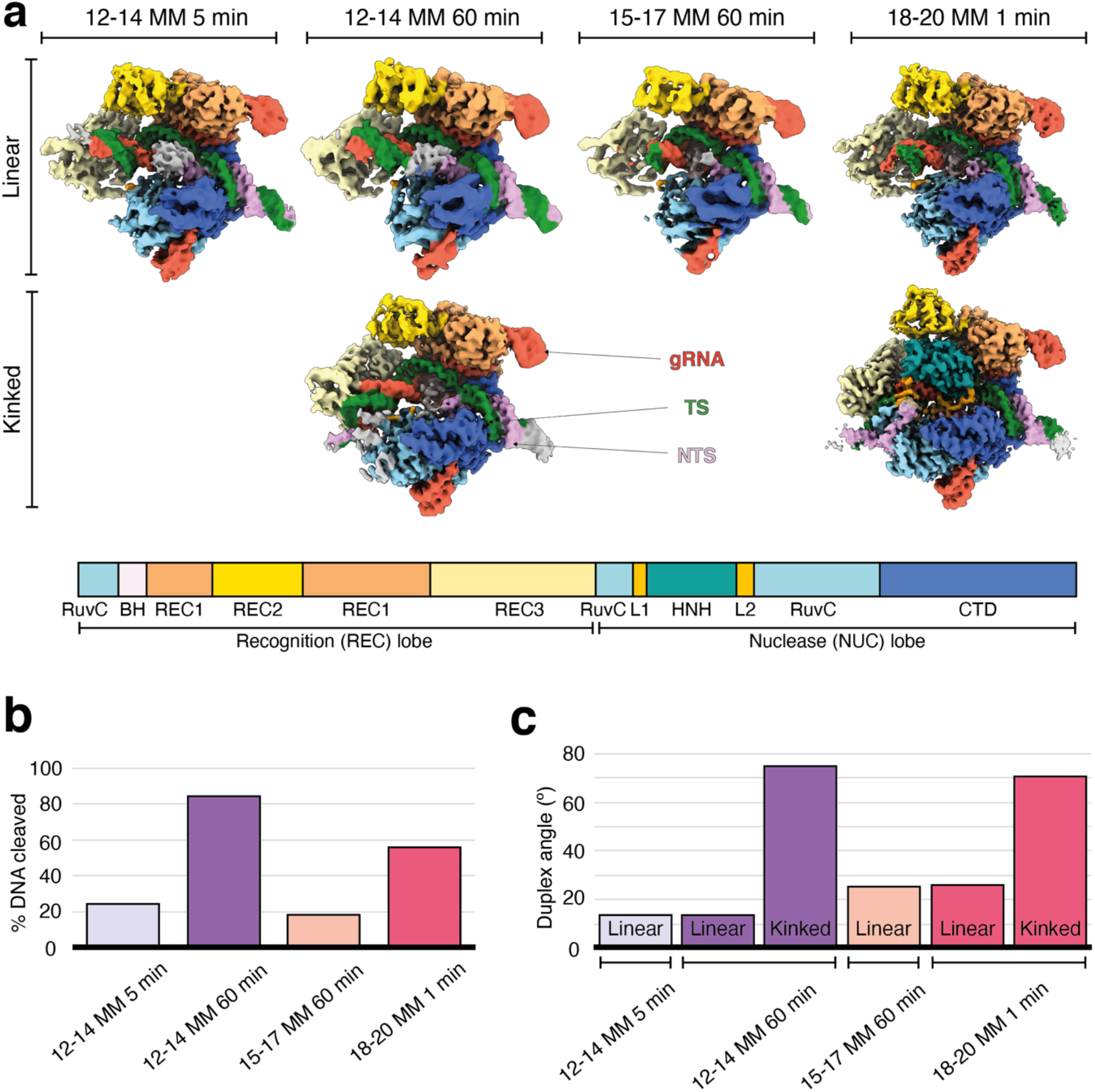
Mismatch-induced Cas9 conformational intermediates. **a**, Cryo-EM reconstructions of Cas9 in complex with various partially mismatched DNA substrates, determined at global resolutions ranging from 2.8-3.6 Å. Cryo-EM maps are colored according to the domain map (below). **b**, Fraction of DNA cleaved by Cas9 at each of the time-pointed used for structure determination. **c**, The between PAM-proximal TS:NTS duplex and PAM-distal gRNA:TS duplex for each of the 6 structures determined.

We then vitrified samples of Cas9 with 12-14 MM DNA after a 60-minute incubation where ∼80% of the DNA was cleaved (**Fig 1b**). Two distinct conformations were observed – a linear duplex conformation consistent with the 5-minute structure of 12-14 MM and the kinked duplex conformation described above (**Fig 1a,c**). The Cas9 conformations in these two structures are identical (**Extended Data Fig. 3a**), but the PAM-distal gRNA:TS duplex end has stably docked with REC3. We propose that the linear duplex conformation corresponds to an early structural intermediate of Cas9, prior to HNH rearrangement and docking to cleave the DNA (Chen et al., 2017; Zhu et al., 2019). We hypothesized that mismatches that prevent PAM-distal gRNA:TS duplex from docking on REC3 would have severely reduced DNA cleavage.

To test this hypothesis, we determined a structure of Cas9 with 15-17 MM dsDNA substrate after 60 minutes of incubation with the enzyme (**Fig 1b**). This mismatch inhibits cleavage by Cas9, but still permits DNA binding as measured by high-throughput profiling (Jones et al., 2021) (**Extended Data Fig.4**). We observed only the linear duplex conformation (**Fig 1a,c**). These structures suggest a model whereby a linear duplex conformation precedes the canonical kinked duplex conformation required for activation, and mismatches that block the kinked conformation abrogate DNA cleavage by Cas9.

### 18-20 mismatch induces a fully active Cas9 conformation

We next sought to understand how certain mismatches can evade Cas9 surveillance and facilitate faster Cas9 activation relative to other mismatches. We examined Cas9 after incubation with 18-20 MM DNA at the 1-minute time-point where ∼50% of the DNA has been cleaved to determine if this more tolerated mismatch undergoes the same structural transition as with 12-14 MM. We observed a mixed population of particles including the linear conformation (**Fig 1a,c**) and the kinked duplex conformation. In the kinked duplex structure, we observed HNH docked at the TS scissile phosphate, indicating the fully active conformation. This arrangement of HNH is entirely consistent with the previously observed active Cas9 conformation (Sternberg et al., 2015; Zhu et al., 2019). Importantly, these results suggest that the population of particles showing a linear conformation represents an early intermediate in the pathway and the kinking of the DNA:RNA duplex is linked to HNH docking.

Consistent with previous studies, we observed TS cleavage between nucleotides 3 and 4 (**Fig 2**) and NTS cleavage at the canonical site 3 bases upstream from PAM. We report the first direct observation of a RuvC active site with the non-target strand bound in a catalytically competent conformation. R986 is in the predicted ‘down’ conformation, allowing it to position a water molecule for nucleophilic attack (**Fig 2**) while F916 wedges between the -2 and -3 bases via stacking interactions, positioning the -3 position within the RuvC active site. This is consistent with previously described molecular dynamics simulations (Casalino et al., 2020).

**Fig. 2.**
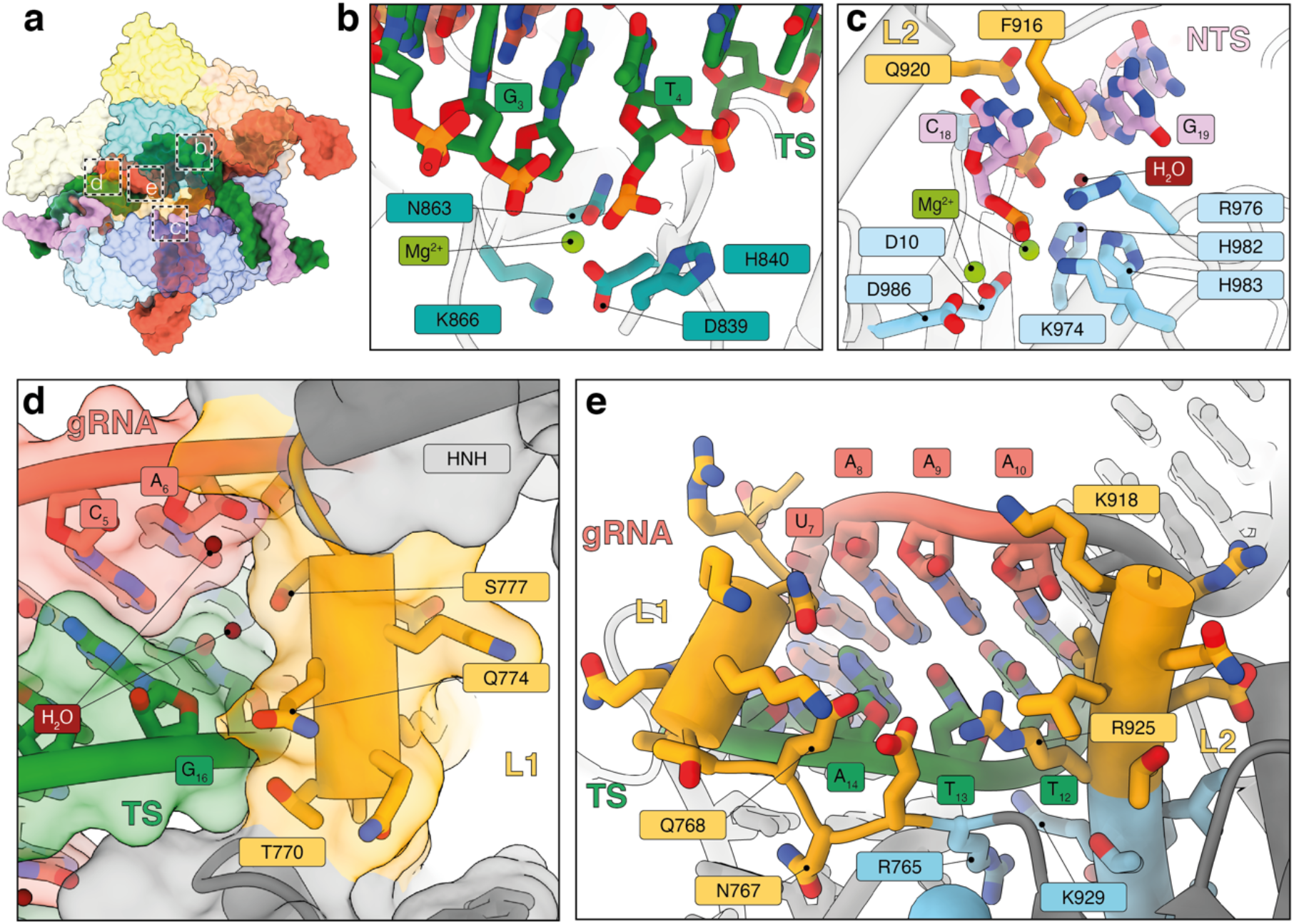
Linkers L1 and L2 mediate structural transition to active state. **a**, Overview of 18-20 MM active conformation. **b,c**, Detailed view of HNH (**b**) and RuvC (**c**) active sites. **d**, Docking of L1 linker helix against PAM-distal gRNA:TS duplex. **e**, Interactions of L1 and L2 regions with minor groove of gRNA:TS duplex. HNH extending from L1 and L2 linkers has been removed for clarity and does not interact with this region of the gRNA:TS duplex.

The fully active configuration requires dramatic conformational rearrangements, including a ∼140° rotation of the HNH domain from the inactive state. Furthermore, our structures reveal the molecular mechanisms underlying this rearrangement. The L1 and L2 linker domains tether HNH to the rest of Cas9 and are often missing from crystal structures, presumably due to their intrinsic flexibility. However, in our active structure, we observe high quality density for both L1 and L2. Notably, L1 helix docks against the minor groove of the PAM-distal gRNA:TS duplex and forms an intricate network of interactions, including multiple water-mediated hydrogen bonds with both strands (**Fig 2**). Since L1 docks on the minor groove these interactions are gRNA:TS structure-specific rather than sequence specific and can only occur when the PAM-distal duplex end is in the kinked conformation. This provides a structural basis for our observation that the kinked duplex conformation is a prerequisite for Cas9 activation and cleavage. Comparisons of our model with Cas9 structures in inactive (EMD-3276) and active (EMD-0584) conformations confirmed that L1 docking against the gRNA:TS duplex is correlated with HNH rearrangement and Cas9 activation (**Extended Data Fig 3**). Furthermore, our observation of L1 and L2 “locking” HNH in an active conformation is supported by the slow rate of dissociation of Cas9 from target DNA post-cleavage (Aldag et al., 2021).

The NTS-stabilizing residue F916 is within the L2 linker domain; however, within the inactive Cas9 conformation, L2 is positioned >20 Å away from the RuvC active site. L1-facilitated positioning of HNH at the TS enables relocation of L2 which in turn enables positioning of the NTS within the RuvC active site. This mechanism provides a structural basis for the observed coupling of TS and NTS cleavage where HNH docking precedes alignment of the NTS at the RuvC site for cleavage (Gong et al., 2018; Liu et al., 2020).

While previous studies have noted the importance of L1 docking onto the gRNA:TS duplex for HNH repositioning (Sun et al., 2019; Zhang et al., 2020), our observation that a linear gRNA:TS duplex conformation induced by PAM-distal mismatches abrogates L1 docking provides a structural explanation for why certain PAM-distal mismatched substrates are able to bind Cas9, while not triggering DNA cleavage (Jones et al., 2021).

### The 18-20 MM is held in a conformation to mimic matching by ordering of a RuvC loop

The 18-20 MM contains an unusual duplex conformation at the site of the mismatch. The C:C mismatch at position 18 on the Target Strand (TS(18)) is stabilized by stacking interactions with adjacent Watson-Crick base pairs. However, the gRNA is otherwise distorted with gRNA-2 flipped out by ∼180° so that gRNA-1 then intercalates between TS(19) & TS(20). TS(19) is then contacted by Q1027:water hydrogen bond, and TS(20) resumes base-pairing with NTS (**Fig 3**).

**Fig. 3.**
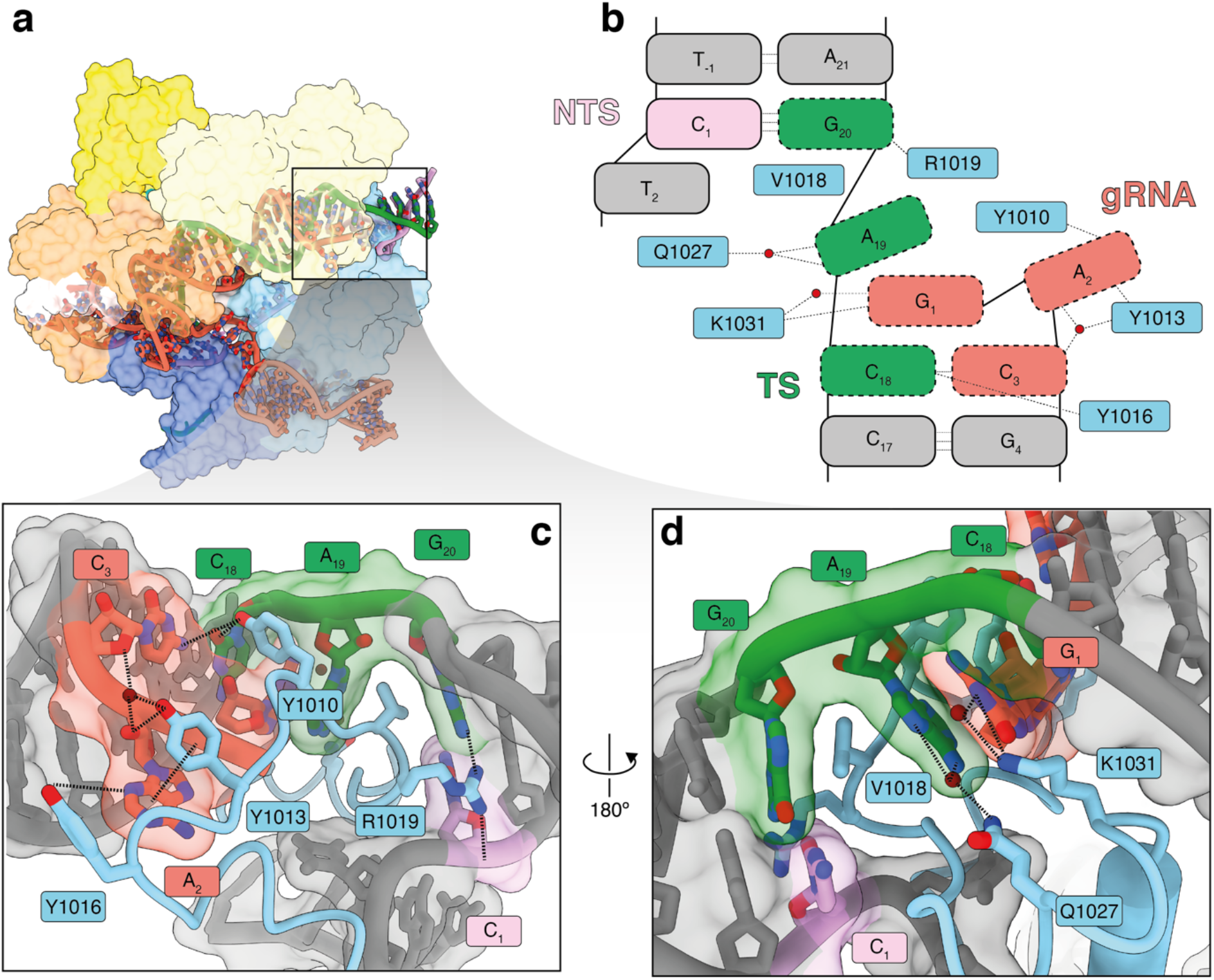
Stabilization of distorted 18-20 MM by RuvC domain. **a**, Overall structure of 18-20 MM active conformation viewed from the back. **b**, Schematic representation of distorted PAM-distal gRNA:TS duplex. Red circles correspond to water molecules. **c & d**, Close-up views of Cas9 interacting with duplex distal end. Flipped gRNA base position 2 is accommodated by stacking interactions and hydrogen bonding with RuvC tyrosine side-chains, while a network of interactions (including a water-mediated hydrogen bond) stabilizes the stretched TS configuration, allowing TS(20) to resume base-pairing with the NTS.

This unusual nucleic acid conformation is stabilized by RuvC and appears to facilitate the binding of this mismatch. The residues within RuvC that contact and stabilize this distorted configuration are absent in previous on-target structures (Anders et al., 2014; Jiang et al., 2016; Zhu et al., 2019), indicating that they are involved only in mismatch binding rather than on-target activation (**Fig 3**). Although this mechanism to facilitate certain mismatches may provide an essential mechanism to reduce mutational escape of Cas9 surveillance by phage, it is counterproductive for use of Cas9 in gene editing.

Previous rationally engineered variants “hyper-accurate Cas9” (HypaCas9: HypaCas9: N692A, M694A, Q695A, and H698A mutations) and “high-fidelity Cas9” (Cas9-HF1: N467A, R661A, Q695A, and Q926A mutations) achieve somewhat higher fidelity at the expense of up to 100-fold reduced efficiency of on-target DNA cleavage (Liu et al., 2020). The mutated residues are mainly located within the REC3 domain and make numerous interactions only with the kinked duplex end. Therefore, by abolishing interactions between REC3 and the PAM-distal duplex, these high-fidelity variants reduce the capacity of Cas9 to stabilize the kinked duplex configuration required for docking of L1 that facilitates HNH repositioning and cleavage activity. Our data provides a structural explanation for why these high-fidelity Cas9 variants prevent activation of Cas9 by off-target substrates, but also act as a throttle for on-target Cas9 activity.

Through kinetics-guided structural determination, we have described a novel gRNA:TS duplex conformational intermediate that precedes Cas9 activation (**Fig 4**). Strikingly, we observe that the well-characterized and widespread off-targeting due to the mismatch at the extreme PAM-distal end (positions 18-20 (Chen et al., 2017; Kuscu et al., 2014; Liu et al., 2020; Sternberg et al., 2015; Tsai et al., 2015)) employs a unique mechanism to for stabilization of a highly distorted duplex conformation, involving a domain loop in RuvC that penetrates the duplex. Excitingly, this region is missing in previously determined structures of Cas9, suggesting that it plays a role solely in mismatch tolerance at these positions. Our results provide molecular insights into the underlying structural mechanisms that govern off-target effects of Cas9 and provide a molecular blueprint for the design of next-generation high fidelity Cas9 variants that reduce off-target DNA cleavage, while retaining efficient cleavage of on-target DNA.

**Fig. 4.**
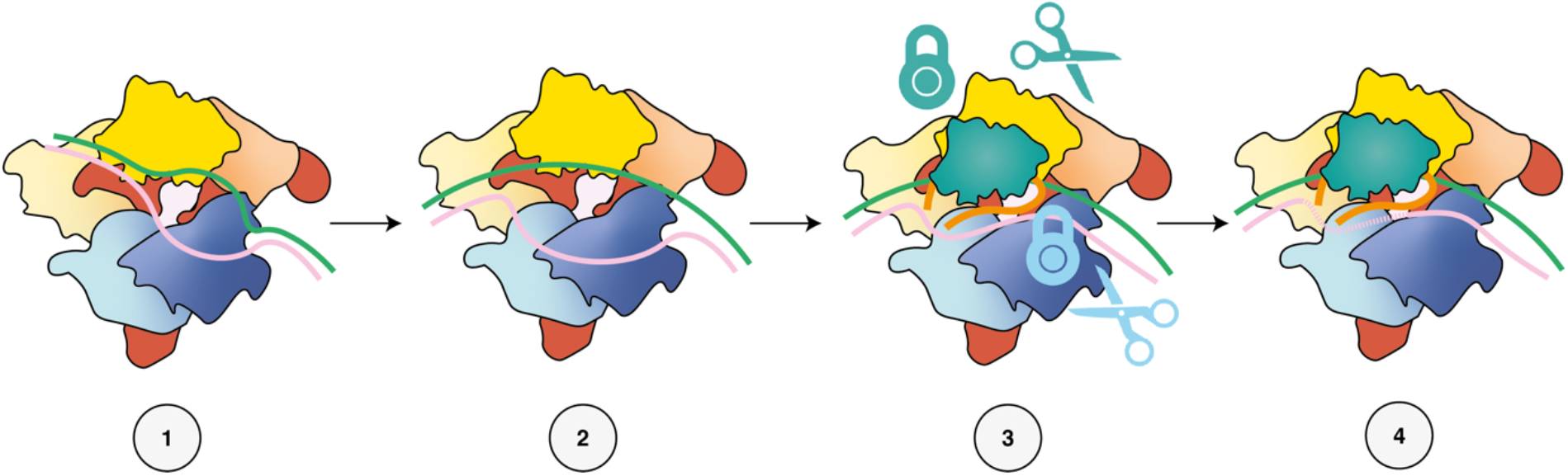
Model for Cas9 activation. At step 1, the gRNA:TS duplex is linear and flexible. Docking of the PAM-distal end of the duplex in the REC3 domain stabilizes the kinked conformation (step 2). Once the kinked conformation has been stabilized, L1 and L2 are able to use the duplex as a structural scaffold and lever HNH to the TS cleavage position. L2 then positions NTS within the RuvC active site. After cleavage, L1 and L2 interactions are maintained with gRNA:TS duplex and NTS, respectively, and HNH remains in place. Mutagenesis of residues within REC3 responsible for PAM-distal end docking destabilize the kinked conformation and therefore prevent L1 docking and Cas9 activation.

## Author Contributions

J.P.K.B. prepared samples for and performed cryo-EM, structure determination, and modeling. M-S.L performed kinetic studies and prepared samples for cryo-EM. K.J. purified SpCas9 and MDCC-Cas9 used for structure determination and kinetic analysis. R.S.M assisted in analysis of the 12-14 MM 5 min structure. J.P.K.B., M-S.L., D.W.T., and K.A.J. analyzed and interpreted the data and wrote the manuscript. D.W.T. and K.A.J. supervised and secured funding for the studies.

## Acknowledgements

This work was supported in part by Welch Foundation grants F-1604 (to K.A.J.) and F-1938 (to D.W.T.), a Robert J. Kleberg, Jr. and Helen C. Kleberg Foundation Medical Research Grant (to D.W.T.). D.W.T is a CPRIT Scholar supported by the Cancer Prevention and Research Institute of Texas (RR160088).

## Declaration of Competing Interests

The authors are inventors on a patent application based on this work. K.A.J. is the President of KinTek, Corp., which provided the chemical-quench flow instruments and the KinTek Explorer software used in this study.

## Materials and Data Availability

The structures of 12-14MM 5 min, 12-14MM 60 min linear and 18-20MM 1 min kinked active have been and their associated atomic coordinates have been deposited into the Electron Microscopy Data Bank (EMDB) and Protein Data Bank (PDB) with accession codes EMD-24833, EMD-24835, EMD-24838 and PDB codes 7S4U, 7S4V and 7S4X, respectively. Maps of 12-14MM 60 min linear, 15-17MM 60 min linear and 18-20 1 min linear have been deposited into the Electron Microscopy Data Bank (EMDB) with accession codes EMD-23834, EMD-24836 and EMD-24837, respectively. All materials are available upon request from David W. Taylor.

## Materials and methods

### Protein expression and purification

SpCas9 was expressed and purified as described previously (Liu et al., 2020).

### Nucleic acid preparation

55-nt DNA duplexes were prepared from unmodified, synthesized DNA oligonucleotide and were PAGE-purified by Integrated DNA Technologies. The DNA duplex used for HNH cleavage assays was prepared using 6-FAM 5’-end labeling of the target strand before annealing with cold non-target strand at a 1:1.15 molar ratio. We purchased the sgRNA for from Synthego. The sequences of the synthesized oligonucleotides including the positions of mismatches are listed in the Table 1.

**Table 1.**
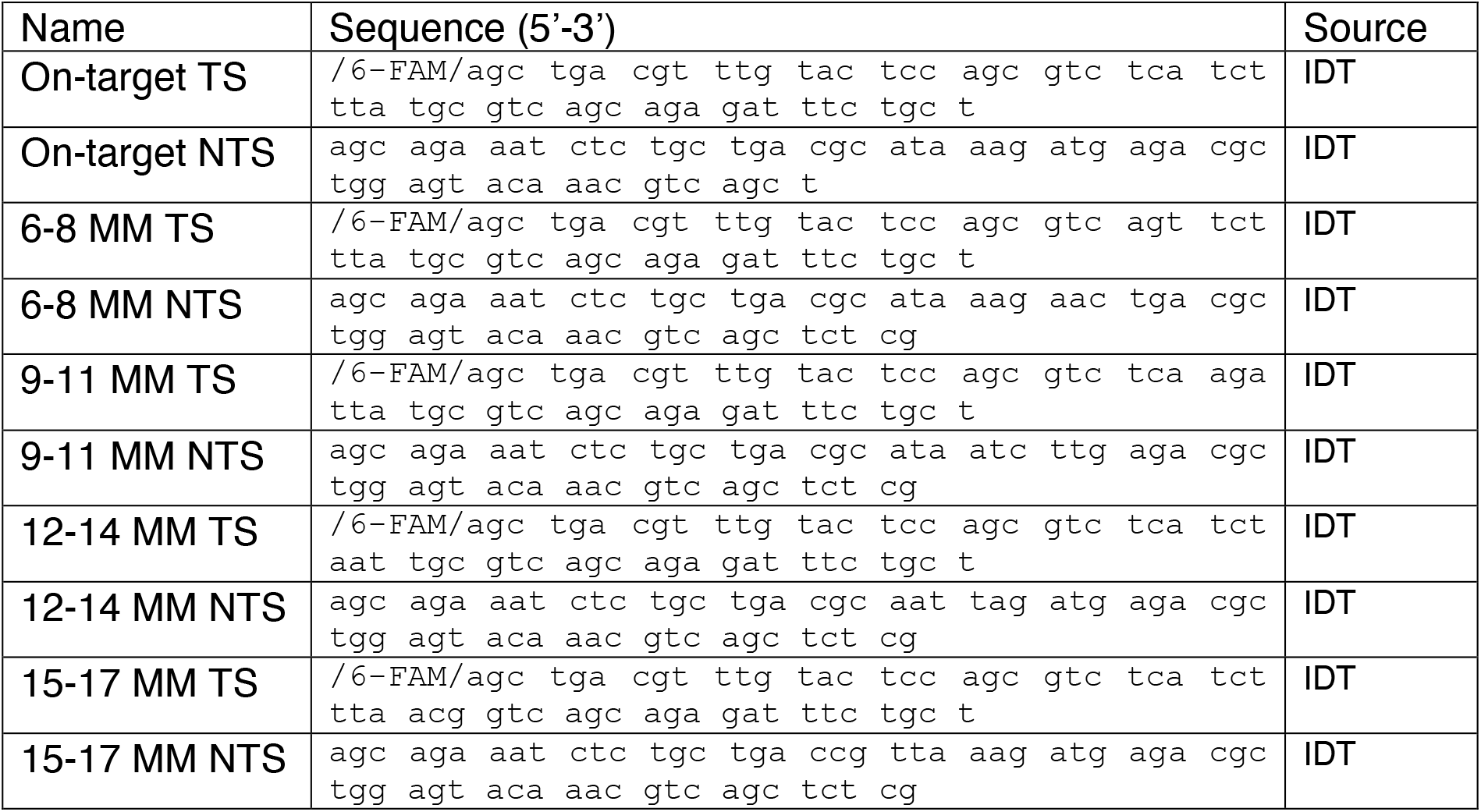
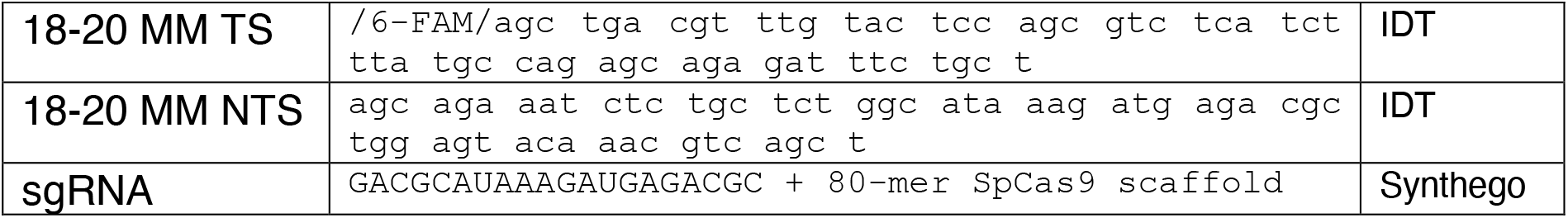
List of nucleotide sequences used in study.

**Table 2.** correlation between fraction DNA cleaved and fraction of cryo-EM particles in linear or kinked duplex conformations.

| Mismatch | Timepoint | % DNA cleaved | % of particles in linear/kinked conformation |
| --- | --- | --- | --- |
| 12-14 | 5 min | 9 | 100/0 |
| 12-14 | 60 min | 82 | 42/58 |
| 15-17 | 60 min | 19 | 100/0 |
| 18-20 | 1 min | 63 | 61/39 |

### Kinetics

#### Buffer composition for kinetic reactions

We prepared 5X cleavage buffer (100 mM Tris-Cl, pH 7.5, 500 mM KCl, 25% glycerol, 5 mM DTT). The time course of DNA cleavage was monitored after mixing Cas9– gRNA with substrate in 1X cleavage buffer (20 mM Tris-Cl, pH 7.5, 100 mM KCl, 5% glycerol, 1 mM DTT).

#### DNA cleavage kinetics

The reaction of Cas9 with on- and off-target was started by mixing Cas9.gRNA (28 nM active-site concentration) with 10 nM fluorescently labeled DNA target in the presence of Mg^2+^ and then stopped at various times by mixing with 0.3 M EDTA (Extended Data Fig. 1). The target strand was labeled with 6-FAM on the 5’-end. Products of the reaction were resolved and quantified using an Applied Biosystems DNA sequencer (ABI 3130xl). Data were fit using either a single or double-exponential equations shown below:

Single exponetial equation:

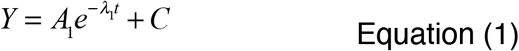

where *Y* represents concentration of cleavage product, *A*_*1*_ represents the amplitude, and *λ*_1_ represents the observed decay rate (eigenvalue).

Double exponetial equation:

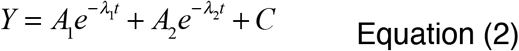

where *Y* represents concentration of cleavage product, *A*_*1*_ represents the amplitude and *λ*_1_ represents the observed rate for the first phase. *A*_*2*_ represents the amplitude and *λ*_2_ represents the observed rate for the second phase.

### CryoEM sample preparation, data collection and processing

Cas9 in complex with various mismatched DNA substrates was frozen at different timepoints, based on kinetic analysis (Extended Data Fig. 1). A non-productive mismatch complex (15-17MM, 1h), a slow productive mismatch (12-14) at early (5 min) and late (1h) time points, and a fast productive mismatch (18-20, 1 min) were chosen. MDCC-Cas9 was used for structure determination in order to couple structural analysis with ongoing kinetic studies monitoring changes in fluorescence. We previously showed the kinetics MDCC-Cas9 were indistinguishable from wild type enzyme (Liu et al., 2020). The cleavage reaction was triggered by mixing 10 *µ*M DNA duplex preincubated with 10 mM MgCl_2_ and 8 *µ*M MDCC-labelled Cas9: 8 *µ*M gRNA was preincubated with 10 mM MgCl_2_, in reaction buffer (19 mM Tris-Cl, pH 7.5, 95 mM KCl, 4.75% glycerol, and 5 mM DTT) at a 1:1 ratio. 4 *µ*l of sample was applied to glow discharged holey carbon grids (C-flat 2/2, Protochips Inc.), blotted for 1 s with a blot force of 4 and rapidly plunged into liquid nitrogen-cooled ethane using an FEI Vitrobot MarkIV. Reactions were quenched through vitrification.

Data were collected on an FEI Titan Krios cryo-electron microscope equipped with a K3 Summit direct electron detector (Gatan, Pleasanton, CA). Images were recorded with SerialEM (Mastronarde, 2005) with a pixel size of 1.1Å for 12-14MM datasets, and 0.81Å for 18-20MM and 15-17MM datasets, over a defocus range of -1.5 to -2.5 *µ*m. During collection of the 12-14MM 5 min timepoint dataset, a preferred orientation was observed. To ameliorate this, a second dataset was collected at 30° tilt. Movies were recorded at 13.3 electrons/pixel/second for 6 seconds (80 frames) to give a total dose of 80 electrons/pixel. CTF correction, motion correction and particle picking were performed in real-time using cryoSPARC live. Further data processing was performed cryoSPARC v3.2 (Punjani et al., 2017).

Multiple rounds of 3D classification within cryoSPARC yielded reconstructions of 6 distinct Cas9 complexes at resolutions ranging from 2.7-3.6Å (**Table 3**). Non-uniform refinement was used for final reconstructions (Punjani et al., 2020). Active Cas9 (PDB ID 6O0X) was rigid-body fitted into each map using ChimeraX (Pettersen et al., 2021). Regions of the model not present in a given map were truncated, and flexible fitting was performed using Namdinator (Kidmose et al., 2019). Further modelling was performed using Isolde (Croll, 2018), and the models were ultimately subjected to real-space refinement as implemented in Phenix.

**Table 3.** Cryo-EM data collection, refinement and validation statistics.

|  | 12-14 MM 5<br>min (linear)<br>(EMD-24833)<br>(PDB 7S4U) | 12-14 MM<br>60 min<br>(linear)<br>(EMD-24834) | 12-14 MM 60<br>min (kinked<br>pre-active)<br>(EMD-24835)<br>(PDB 7S4V) | 15-17 MM<br>60 min<br>(linear)<br>(EMD-24836) | 18-20<br>MM 1<br>min<br>(linear)<br>(EMD-24837) | 18-20 MM 1<br>min (kinked)<br>(EMD-24838)<br>(PDB 7S4X) |
| --- | --- | --- | --- | --- | --- | --- |
| **Data collection and processing** |  |  |  |  |  |  |
| Magnification | 22,500 | 22,500 | 22,500 | 29,000 | 29,000 | 29,000 |
| Voltage (kV) |  |  | 300 kV |  |  |  |
| Electron exposure (e<sup>-</sup>/Å<sup>2</sup>) | 80 | 80 | 80 | 70 | 70 | 70 |
| Defocus range (μm) |  |  | -1.5 to -2.5 |  |  |  |
| Pixel size (Å) | 1.1 | 1.1 | 1.1 | 0.81 | 0.81 | 0.81 |
| Symmetry imposed |  |  | C1 |  |  |  |
| Initial particle images (no.) | 1,546,987 | 1,185,683 | 1,185,683 | 1,200,112 | 997,043 | 997,043 |
| Final particle images (no.) | 376,601 | 139,113 | 198,005 | 222,693 | 163,647 | 104,658 |
| Map resolution (Å) | 3.57 | 3.47 | 3.28 | 3.33 | 2.87 | 2.76 |
| FSC threshold |  |  |  |  |  |  |
| Map resolution range (Å) |  | 3 - 7 |  |  |  | 2 - 6 |
| **Refinement** |  |  |  |  |  |  |
| Initial model used (PDB code) | 6O0X | N/A | 6O0X | N/A | N/A | 6O0X |
| Model resolution (Å) | 4.2 | N/A | 3.4 | N/A | N/A | 3.0 |
| FSC threshold | 0.5 | 0.5 | 0.5 |  |  | 0.5 |
| Map sharpening *B* factor (Å<sup>2</sup>) | 202.1 | 92.0 | 96.8 | 111.7 | 59.3 | 68.8 |
| Model composition |  |  |  |  |  |  |
| Non-hydrogen atoms | 12099 | N/A | 12221 | N/A | N/A | 14511 |
| Protein residues | 1092 |  | 1122 |  |  | 1354 |
| Nucleotides | 150 |  | 145 |  |  | 161 |
| Ligands | 0 |  | 0 |  |  | 0 |
| Mean *B* factors (Å<sup>2</sup>) |  |  |  |  |  |  |
| Protein | 93.68 | N/A | 30.98 | N/A | N/A | 42.14 |
| Nucleotides | 80.32 |  | 53.58 |  |  | 75.72 |
| R.m.s. deviations |  |  |  |  |  |  |
| Bond lengths (Å) | 0.013 | N/A | 0.004 | N/A | N/A | 0.002 |
| Bond angles (°) | 2.017 |  | 0.555 |  |  | 0.507 |
| Validation |  |  |  |  |  |  |
| MolProbity score | 1.28 | N/A | 1.58 | N/A | N/A | 1.59 |
| Clashscore | 0.49 |  | 4.66 |  |  | 6.62 |
| Poor rotamers (%) | 1.84 |  | 0 |  |  | 0 |
| Ramachandran plot |  |  |  |  |  |  |
| Favored (%) | 93.87 | N/A | 95.05 | N/A | N/A | 96.81 |
| Allowed (%) | 5.95 |  | 4.95 |  |  | 3.12 |
| Disallowed (%) | 0.19 |  | 0 |  |  | 0.07 |

## Supplementary materials

**Extended Data Fig. 1.**
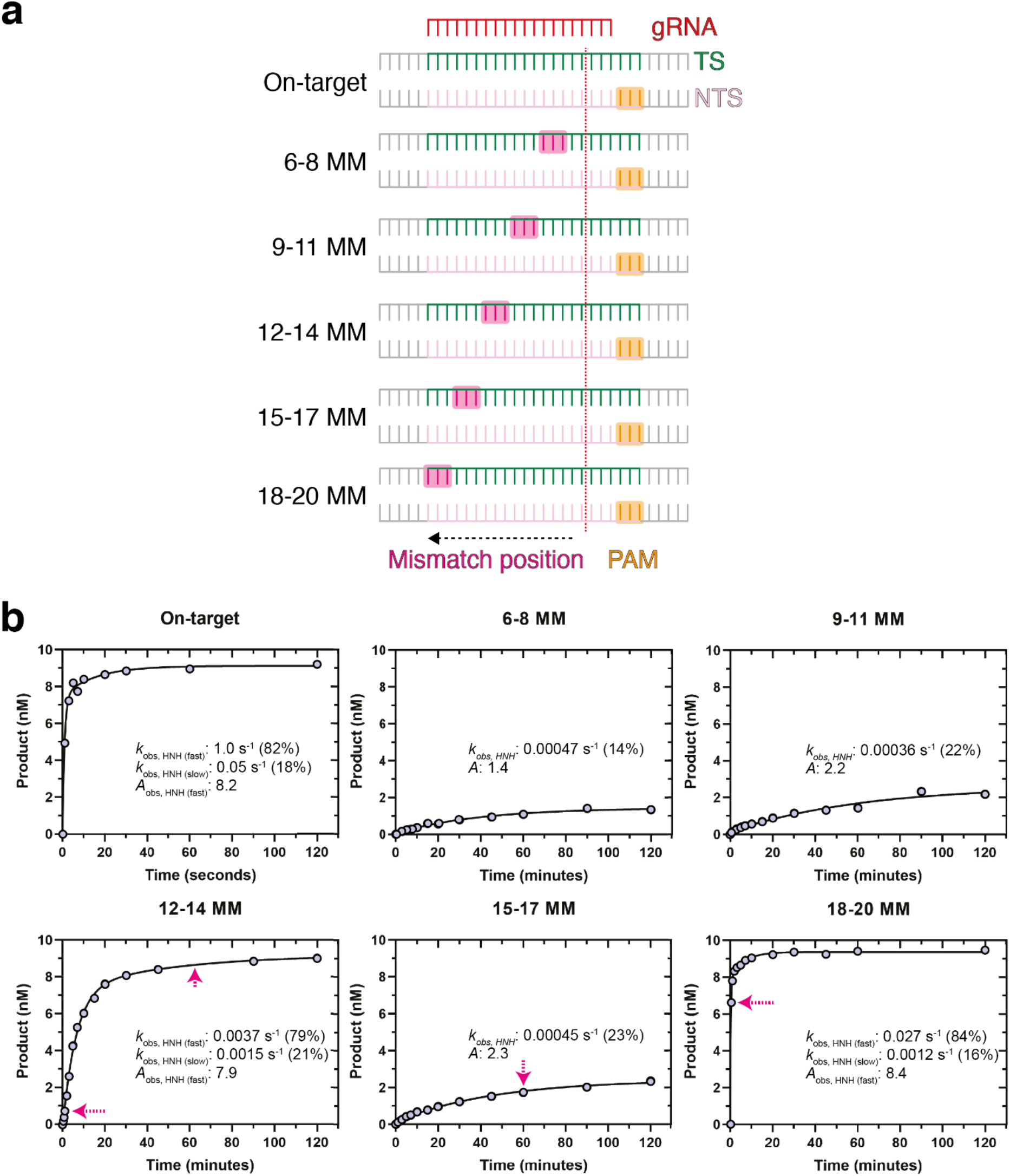
Kinetic basis for mismatch surveillance by Cas9. **a**, Schematic representation of mismatch scanning constructs used for kinetic analysis. **b**, Time course of cleavage of on-target and mismatched DNA (10 nM) by Cas9. Magenta arrows correspond to time-points used to prepare cryo-EM samples.

**Extended Data Fig. 2.**
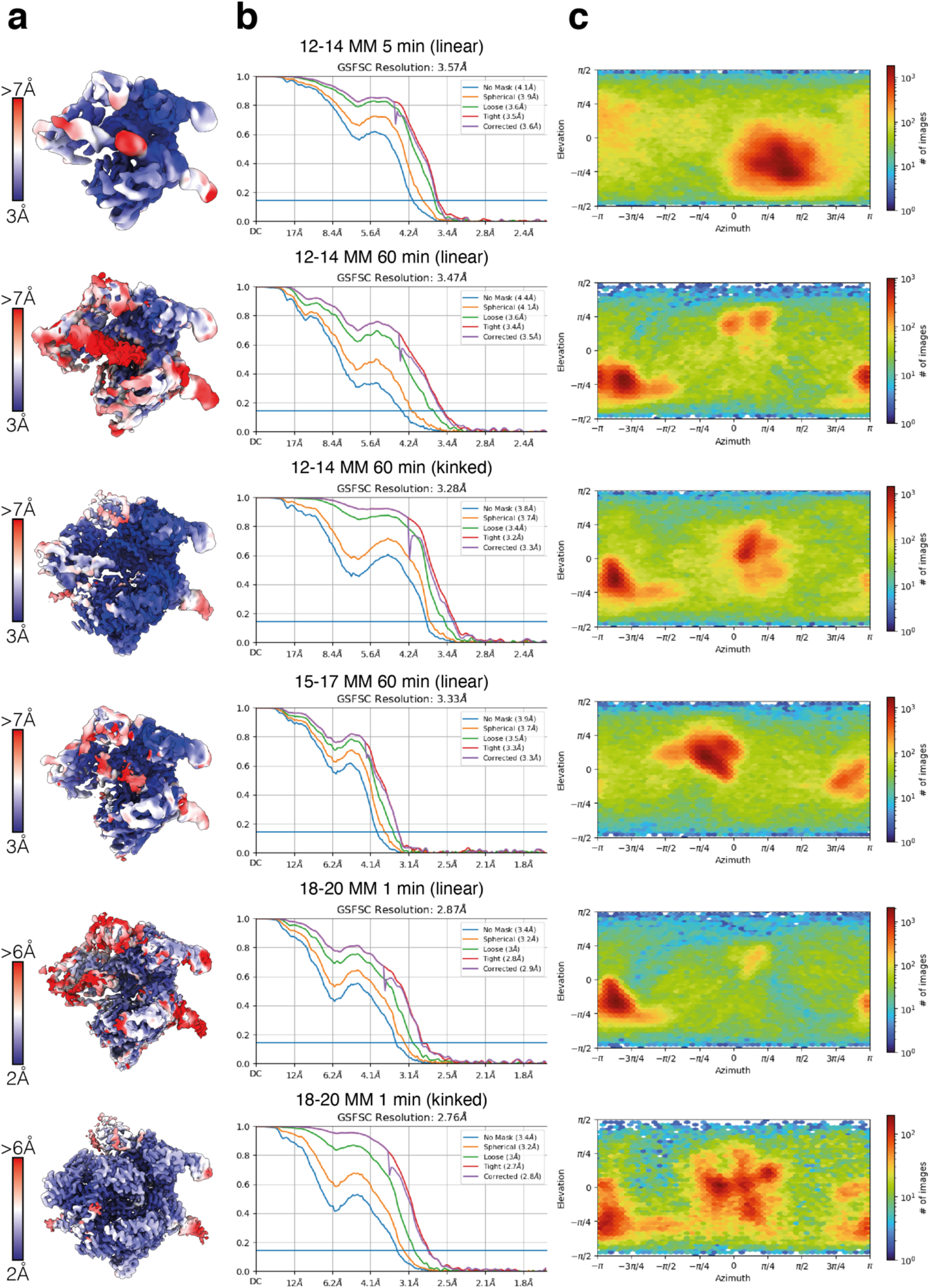
Resolution estimates and orientation distributions of cryo-EM maps. **a**, Unsharpened maps colored according to local resolution. **b**, Gold-standard FSC curves for cryo-EM reconstructions. Resolutions were estimated at FSC=0.143. **c**, Euler diagrams showing orientation distributions of cryo-EM reconstructions.

**Extended Data Fig. 3.**
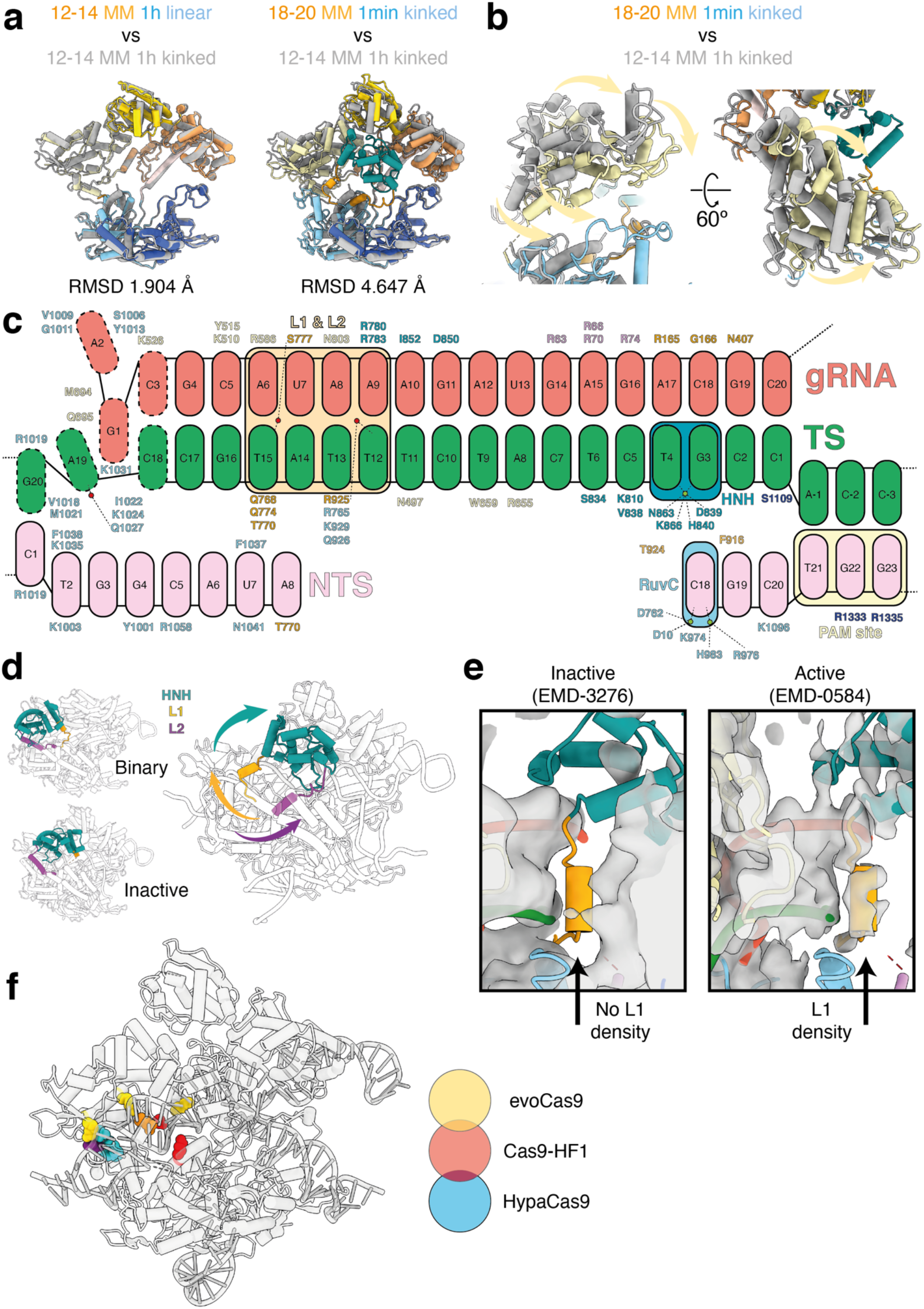
Structural analysis of Cas9. **a**, Left: Comparison of Cas9 only 12-14 MM 60 min linear (color) and only 12-14 MM 60 min kinked (grey) models. Right: Comparison of Cas9 only active conformation (18-20 MM 1 min linear, color) and kinked pre-active (12-14 MM 60 min kinked, grey) models. While there is no significant conformational change associated between transition from linear to kinked pre-active (root-mean standard deviation (RMSD) between equivalent Cα atoms of 1.904 Å), the change from kinked pre-active to active conformations is associated with a larger conformational change (4.647 Å, most of which occurs within the REC3 domain). **b**, Close-up view of REC3 conformational changes that occur upon activation, as viewed from the left side. REC3 moves forwards towards the kinked duplex by up to almost 15 Å upon activation and HNH repositioning. **c**, Schematic representation of Cas9 : nucleic acid contacts. **d**, Conformations of HNH domain (green) and L1 (gold) and L2 (purple) linkers in the context of Cas9 binary complex (i.e. with gRNA, PDB 4ZT0), Cas9:gRNA complex bound to dsDNA in an inactive conformation (PDB 5F9R), and in the active Cas9 18-20 MM structure presented in this work. Upon activation, HNH is repositioned at the TS cleavage site, driven by large conformational changes in the L1 and L2 linkers. **e**, Comparison with the active Cas9 18-20 MM structure presented in this work and previously determined cryo-EM maps (transparent grey) of inactive (left, EMD-3276 (Jiang et al., 2016)) and active (right, EMD-0584 (Zhu et al., 2019)) Cas9 bound to on-target dsDNA. The inactive Cas9 has no density for L1 helix at the kinked distal-docked gRNA:TS site, whereas there is clear density for L1 at this site in the active Cas9 cryo-EM map. **f**, Mapping of residues mutated to alanine in selected high-fidelity Cas9 variants. EvoCas9 (yellow) – M495, Y515, R661, K526. Cas9-HF1 (red) – N497, R661, Q695, Q926. HypaCas9 (blue) – N692, M694, Q695, H698. Residues shared between Cas9-HF-1 and either EvoCas9 or HypaCas9 are shown as orange and purple, respectively.

**Extended Data Fig. 4.**
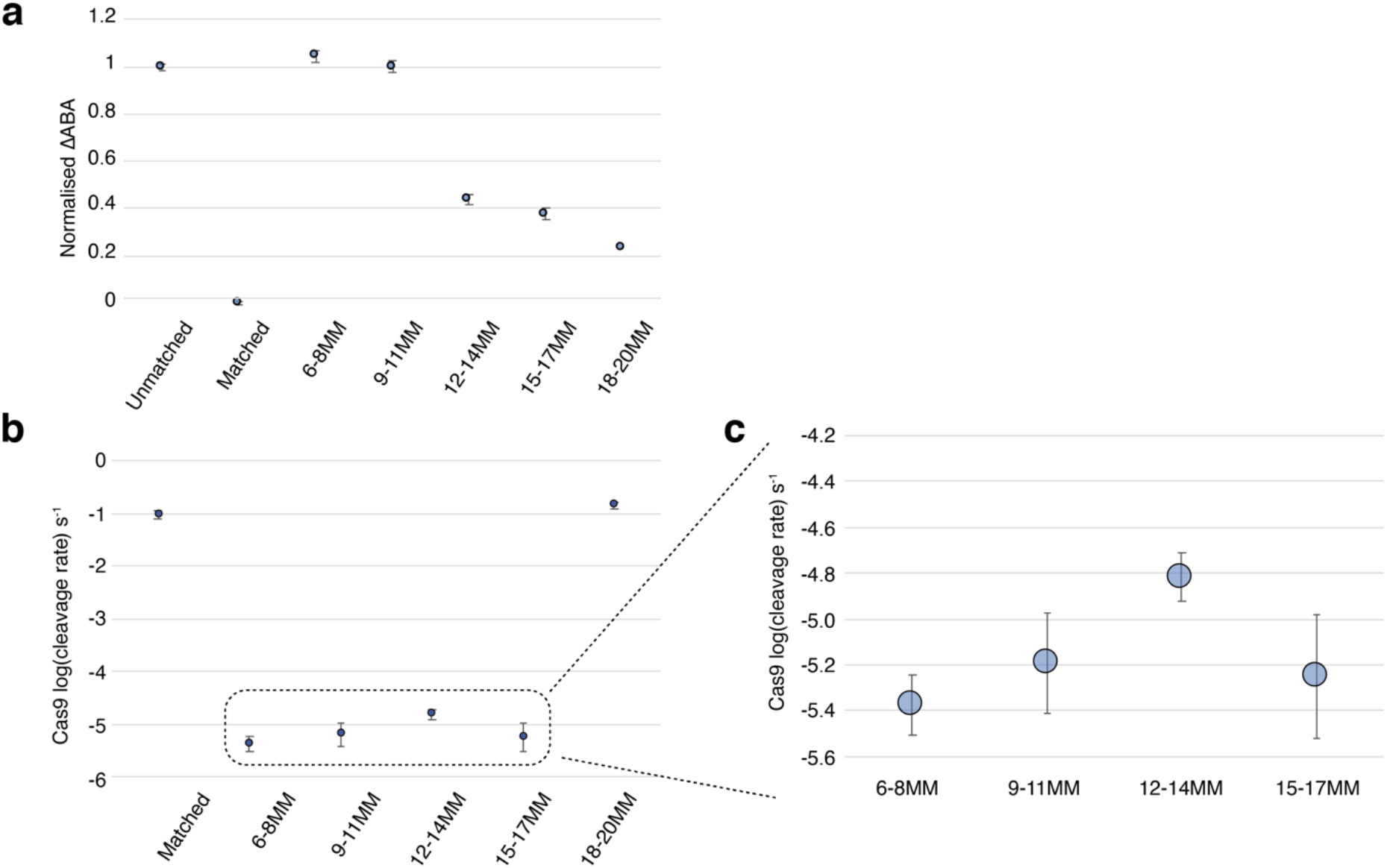
High-throughput analysis of Cas9 DNA binding and cleavage. **a**, Deactivated Cas9 (dCas9) change in apparent binding affinities (ABA) for fully unmatched, fully complementary (matched), and triple contiguous mismatches as shown in Extended Data Fig. 1. ABAs are normalized to matched and unmatched targets. 6-8 MM and 9-11 MM show poor binding to dCas9, whereas 12-14 MM, 15-17 MM and 18-20 MM have binding affinities closer to fully complementary sequences. **b**, DNA cleavage rates by Cas9. **c**, Zoom-in of 6-8 MM, 9-11 MM, 12-14 MM and 15 - 17 MM. 12-14 MM is cleaved 2.5 – 3.5-fold faster than the other triple mismatches presented in this panel.

## Notes

### Competing Interest Statement

The authors are inventors on a submitted patent application based on this work. K.A.J. is the President of KinTek, Corp., which provided the chemical-quench flow instruments and the KinTek Explorer software used in this study.

